# Continual reelin signaling by the prime neurogenic niche of the adult brain

**DOI:** 10.1101/2021.12.12.472284

**Authors:** F. Javier Pérez-Martínez, Manuel Cifuentes, Juan M. Luque

## Abstract

During development reelin sets the pace of neocortical neurogenesis enabling in turn newborn neurons to migrate, but whether and, if so, how reelin signaling affects the adult neurogenic niches remains uncertain. We show that reelin signaling, resulting in Dab1 phosphorylation, occurs in the ependymal-subependymal zone (EZ/SEZ) of the lateral ventricles where, along with its associated rostral migratory stream (RMS), the highest density of functional ApoER2 accumulates. Mice deficient for reelin, ApoER2 or Dab1 exhibit enlarged ventricles and dysplastic RMS. Moreover, while the conditional ablation of Dab1 in neural progenitor cells (NPCs) enlarges the ventricles and impairs neuroblasts clearance from the SEZ, the transgenic misexpression of reelin in NPCs of reelin-deficient mice normalizes the ventricular lumen and the density of ependymal cilia, ameliorating in turn neuroblasts migration; consistently, intraventricular infusion of reelin reroutes neuroblasts. These results demonstrate that reelin signaling persists sustaining the germinal niche of the lateral ventricles and influencing neuroblasts migration in the adult brain.

## Introduction

Neurogenesis in the embryonic and adult stages emerges like a continuous event as the lineage of neural progenitor cells (NPCs) and the conservation of many intrinsic signaling pathways are being revealed (reviewed by Kriegstein and Alvarez-Buylla, 2009; Ming and Song, 2011). The origin and characteristics of extrinsic signals might still challenge this notion.

Radial glia during development and a subpopulation of radial glia-derived like-astrocytes in the adult brain are regarded as the founder cells for most, if not all, neural lineages in the central nervous system (CNS). These neural stem cells undergo distinct modes of cell division, giving rise to the diversity of macroglial and neuronal cell types in the CNS (reviewed by Kriegstein and Alvarez-Buylla, 2009). Particularly, neural cells do not function in their birth sites but undergo more or less extensive migrations. Active adult neurogenesis largely occurs in two restricted regions of the forebrain, the ependymal-subependymal zone (EZ/SEZ) of the lateral ventricles and the subgranular zone (SGZ) in the dentate gyrus of the hippocampus. The EZ/SEZ in the wall of the lateral ventricles constitutes the largest neurogenic niche of the adult mammalian brain (reviewed by Ihrie and Alvarez-Buylla, 2011). It contains multiple cell populations including ependymocytes lining the ventricle and SEZ-astrocytes that are thought to behave as neural stem cells (so-called type B cells). Type B cells give rise to actively proliferating C cells that function as intermediate progenitor cells. C cells in turn generate immature neuroblasts (type A cells) which migrate in chains through the rostral migratory stream (RMS) to the olfactory bulb (OB) where they differentiate into granular and periglomerular interneurons (Lois and Alvarez-Buylla, 1994; Doetsch et al., 1999; Carleton et al., 2003; Brill et al., 2009). Type B and C cells are also closely associated to blood vessels and to a nearby large extracellular matrix (Mercier et al., 2002; Shen et al., 2008; Tavazoie et al., 2008). Type B cells retain important properties of radial glial cells, considered in turn neural stem cells during brain development. For example, their cell bodies are generally located just under the EZ cell layer but have short processes that extend through the ependyma with small apical endings on the ventricle contacting the cerebrospinal fluid (CSF) (Mirzadeh et al., 2008; Shen et al., 2008). Thus, the SEZ can be regarded as a sort of displaced neuroepithelium. Interestingly, neuroblasts migration parallels CSF flow and beating of ependymal cilia is required for normal CSF flow, concentration gradient formation of CSF molecules, and directional migration of neuroblasts (Sawamoto et al., 2006).

The conservation of many intrinsic signaling pathways underlies the remarkable similarity between embryonic and adult neurogenesis, although the origin and nature of extrinsic signals might be different (reviewed by Ming and Song, 2011). A range of morphogens, including Notch, many Ephrins and Eph receptors, growth factors, neurotrophis, cytokines, neurotransmitters and hormones have been identified as major extracellular players of adult neurogenesis (reviewed by Ming and Song, 2011; Zhao et al., 2008; Jang et al., 2008). A rather limited knowledge exists of extracellular cues controlling targeted neuronal migration during adult neurogenesis, or about the molecular links connecting neurogenesis and migration. A few adhesion molecules (such as β1-integrin, PSA-NCAM and tenascin-R) and extracellular cues (such as neuregulins and slits) are known to regulate the stability, motility, or directionality of neuronal migration during EZ/SEZ neurogenesis (reviewed by Lledo et al., 2006; Ming and Song, 2005).

The secretable glycoprotein reelin and its signaling machinery are well known to be required for proper brain development, affecting neuronal migration and positioning of several neuronal lineages. It is well established that reelin binding to Apolipoprotein E Receptor 2 (ApoER2) or Very Low Density Lipoprotein Receptor (Vldlr) induces phosphorylation of the cytoplasmic adaptor protein Disabled 1 (Dab1), a tyrosine kinase signal transduction cascade, and Dab1-regulated turnover. Indeed, the *reeler* mice (deficient in reelin) and the mutant null mice for Dab1 or for both ApoER2 and Vldlr show comparable positioning defects (reviewed by Rice and Curran, 2001; Tissir and Goffinet, 2003; Luque, 2007; Cooper, 2008). However, reelin does not function simply as a positional signal. Rather, it appears to participate in multiple events critical for neuronal migration and cell positioning (Magdaleno et al., 2002). Recently, we have shown that during forebrain development reelin, acting upstream of Notch signaling, it is both required and sufficient to determine the rate of neocortical neurogenesis (Lakoma et al., 2011). Such interaction is also required for neuronal migration and positioning (Hashimoto-Torii et al., 2008). Thus, the early function of reelin in the proliferative compartment might underlie its post-proliferative requirement raising the possibility that acts coupling neurogenesis and neuronal migration, perhaps akin to a permissive rather than instructive signal for cellular migration. Indeed, a reelin-dependent ApoER2 downregulation mechanism uncouples neocortical newborn neurons from NPCs, thereby enabling neurons to migrate (Perez-Martinez et al., 2012). Highlighting the difficulty to establish a coherent model of reelin function, a number of studies has provided exceedingly contradictory evidence as to whether or not reelin plays a role during adult neurogenesis and cell migration (Kim et al., 2002; Hack et al., 2002; Won et al., 2006; Zhao et al., 2007; Heinrich et al., 2006; Andrade et al., 2007; Gong et al., 2007; Simo et al., 2007; Blake et al., 2008; Massalini et al., 2009; Pujadas et al., 2010; Courtes et al., 2011; Teixeira et al., 2012).

In the present study, we use both loss- and gain-of-function genetic approaches along with *in vivo* and *ex vivo* assays to investigate reelin signaling in the adult brain. We seek to reveal whether it persists by the germinal niche of the lateral ventricles. Ours results show that reelin signaling remains active inside, and is necessary and sufficient to modulate, the EZ/SEZ neurogenic niche. They suggest that cerebrospinal fluid reelin acting upstream of ApoER2 and Dab1 regulates the integrity and functionality of the ventricular ependyma, the nearby NPCs, and eventually the differentiation and migratory behavior of neuroblast through the RMS.

## Material and Methods

### Mice

Heterozygous *reeler* mice were purchased from Jackson Laboratory (Bar Harbor, ME). *ApoER2*-null and *Vldlr*-null mice were a gift from Johannes Nimpf (University of Wien, Austria) and were genotyped as described previously (Trommsdorff et al., 1999). *Nestin-reelin* mice were a gift from Tom Curran (The Children’s Hospital of Philadelphia, PA) and were genotyped as previously described (Magdaleno et al., 2002). *Nestin^CreERT2^* mice were a gift from Gordon Fishell (University of New York, NY) and were genotyped as previously described (Balordi and Fishell, 2007). *Dab1*-null mice and conditional (floxed) *Dab1* (*Dab1^c/c^*) mice were obtained and genotyped as described elsewhere (Howell et al., 1997b; Pramatorova et al., 2008). Mutant mice (*Nestin^CreER/+^;Dab1^c/c^*) were generated by crossing homozygous conditional *Dab1* mice (*Dab1^c/c^*) with mice homozygous for the conditional *Dab1* allele and heterozygous for the *Nestin^CreER^* allele (*Nestin^CreER/+^;Dab1^c/c^*). All primers sequences, PCR conditions and the expected sizes of PCR products are available upon request. Animals were handled according to protocols approved by the European Union, NIH guidelines and the Instituto de Neurociencias Animal Care and Use Committees.

### Tamoxyfen and bromodeoxyuridine administration

Tamoxyfen (Sigma, St. Louis, MO) was prepared as a 25 mg/ml stock solution in corn oil (C-8267; Sigma). For Cre induction and analysis of P60 *Nestin^CreER/+^;Dab1^c/c^* mice, tamoxyfen was administrated by intraperitoneal injection (5mg/35g of body weight) every other day for a total of eight infections. After a break of 1 week, another round of eight injections was administered. The recombination effectiveness of this dosage in *Nestin^CreERT2^* mice and the lack of appreciable short or long-effect of tamoxifen exposure on proliferation in the E/SEZ have been previously shown (Balordi and Fishell, 2007). Bromodeoxyuridine (BrdU; Sigma) was administrated intraperitoneally at 100 mg/g of body weight.

### Alkaline phosphatase-reelin *in situ* staining

An alkaline phosphatase (AP)-fusion probe of the receptor-binding region of reelin (repeats 3 to 6, AP-RR36) was generated as previously described (Uchida et al., 2009). Validation of the probe, binding conditions and alternative staining procedures has been also described (Uchida et al., 2009; Perez-Martinez et al., 2012). AP-RR36 *in situ* binding was detected using AP substrates or by double immunofluorescence labeling of AP along with several cell markers.

### Immunohistochemistry

Immunostaining was performed as previously described (Lakoma et al., 2011). Primary and secondary antibodies were acquired and diluted as indicated. Primary antibodies: rabbit anti-human placental alkaline phosphatase (Abcam, 1:100), mouse anti-human placental alkaline phosphatase (Abcam, 1:50), goat anti-doublecortin (Santa Cruz, 1:100), rabbit anti-GFAP (Dako, 1:500), rabbit anti-tyrosine hydroxylase (Pel-Freez, 1:1000), rabbit anti-calbindin (Swant, 1:10000), rat anti-BrdU (Serotec, 1:100), mouse anti-reelin (G10, Abcam, 1:1000), rabbit anti-S100b (Dako, 1:400), rabbit anti-Dab1 (B3, 1:500, [Howell et al., 1997a; Luque et al., 2003]), mouse anti-CRE (Novagen, 1:500), mouse anti-Mash1 (BD Pharmigen, 1:500), mouse anti-phosphotyrosine (Sigma, 1:2000). Secondary antibodies: biotinilated-donkey anti-rabbit, biotinilated-goat anti-rat, biotinilated-donkey anti-goat (Jackson Immunoresearch, 1:500), Cy3-donkey anti-rabbit, Cy3-Goat anti-rat, Cy3-donkey anti-goat, Cy3-donkey anti-mouse, Cy2-donkey anti-rabbit, Cy2-donkey anti-mouse (Jackson Immunoresearch, 1:200) and HRP-goat anti-rabbit, HRP-goat anti-mouse (Jackson Immuno-research, 1:5000). Images were captured on a Leica TCS SP2 AOBS inverted Laser Scanning Confocal Microscope. Image J (NIH, http://rsb.info.nih.gov/ij/), and Adobe Photoshop software were used for image capturing and analysis.

### Production of reelin

Recombinant reelin was produced using the mammalian expression construct pCrl, encoding mouse reelin cDNA (D’Arcangelo et al., 1997), Cell culture and transfection procedures have been previously described (Saez-Valero et al., 2003).

### Dab1 phosphorylation assay

Brains were harvested from P60 *reeler* mice. Two-mm thick coronal slices were obtained and the E/SEZ at the striatal side of the lateral ventricles was microdissected under a low-power dissecting microscope in ice-cold DMEM. Tissues were homogenized in cell lysis buffer and precleared by short centrifugation. Anti-Dab1 immunoprecipitation was carried out from equal amounts of protein, and Dab1 protein levels and phosphorylation status upon reelin stimulation were determined as previously described (Howell et al., 1999).

### Ultrastructural analysis

For scanning electron microscopy, P60 wild type, *reeler* and *reeler ne-reelin* mice were used. Briefly, the entire wall of the lateral ventricles was dissected as whole mount, and fixed with 2.5% glutaraldehyde/2% paraformaldehyde overnight. The tissues were incubated in 2% OsO_4_ for 1h, dehydrated with ethanol, and dried with a critical point drier. After sputter coating with platinum, the mounted samples were examined with a Hitachi S-4100 scanning electron microscope.

### Intraventricular infusion of reelin

Recombinant reelin was produced and concentrated as previously described (Saez-Valero et al., 2003). The concentrated protein (or mock) was infused using a mini osmotic pump (Alzet model 2001) into the caudal region of the lateral ventricle of P60 wild type and *reeler* mice at the coordinate-0.7mm, 1.2mm, 2.0mm (anterior, lateral, depth coordinates relative to bregma). Five days later the brains were processed for immunohistochemistry using anti-doublecortin antibodies.

## Results

### The EZ/SEZ and RMS contain the highest density of functional reelin receptors in the adult brain

To test whether or not the reelin pathway continues to be active in postnatal ages, it was essential to know if there was a continuity in the expression of its functional receptors in adult animals. In fact, we realized that during embryonic development, the lateral ganglionic eminence (LGE), but not the medial ganglionic eminence (MGE), the main origin of the EZ/ SVZ/RMS system of the adult brain (Wichterle et al., 1999, 2001), expresses FRR (**Supplementary Fig. 1**).For this reason, we carried out an analysis of its expression in the adult brain using fusion protein AP-RR36 that we previously developed and extensively validated in isolated cells and embryonic tissues (Uchida et al., 2009; Pérez-Martínez et al., 2012). The excellent resolution offered by the enzymatic development of this technique made it possible to elaborate a rostral-caudal cartography of the expression of FRR, from the olfactory bulb to the brainstem, including the cerebellum (**Supplementary Fig. 2**). Thus, we clearly observed that the highest density of FRR in the adult brain in the two known adult neurogenic niches, the ependymal/subependimal zone (EZ/SVZ) of the lateral ventricles, including the rostral migratory stream (RMS), and the subgranular zone (SGZ) of the dentate gyrus in the hippocampus. A diffuse staining was also observed in the glomerular layer of the olfactory bulb, the medial habenular nucleus and the molecular layer of the cerebellum. Moreover, the greatest contribution to this signal in the neurogenic niches of the brain is due to ApoER2, as is clear when analyzing the null mutants of this receptor, practically lacking staining except for a diffuse signal in the glomerular layer of the olfactory bulb. Consistent with these results are those obtained with the null mutants for the other canonic reelin receptor, VLDLR: the wild-type-like staining of FRR throughout the brain but minimal in the cerebellum (expression practically complementary to that of the mutant ApoER2), allows confirm that VLDLR is the main receptor present in this structure. On the other hand, the staining of the AP-reelin probe shows a gradual increase as the genetic load of reelin decreases. Thus, the signal observed in reeler brain is greater than in the heterocygotic brain for reelin that, in turn, is greater than in the wild-type brain, which supports that, in effect, reelin causes a down-regulation of the receptor. Details of the localization of functional reelin receptors in the adult brain neurogenic niches can be observed in **Figure 1**.

**Fig. 1:**
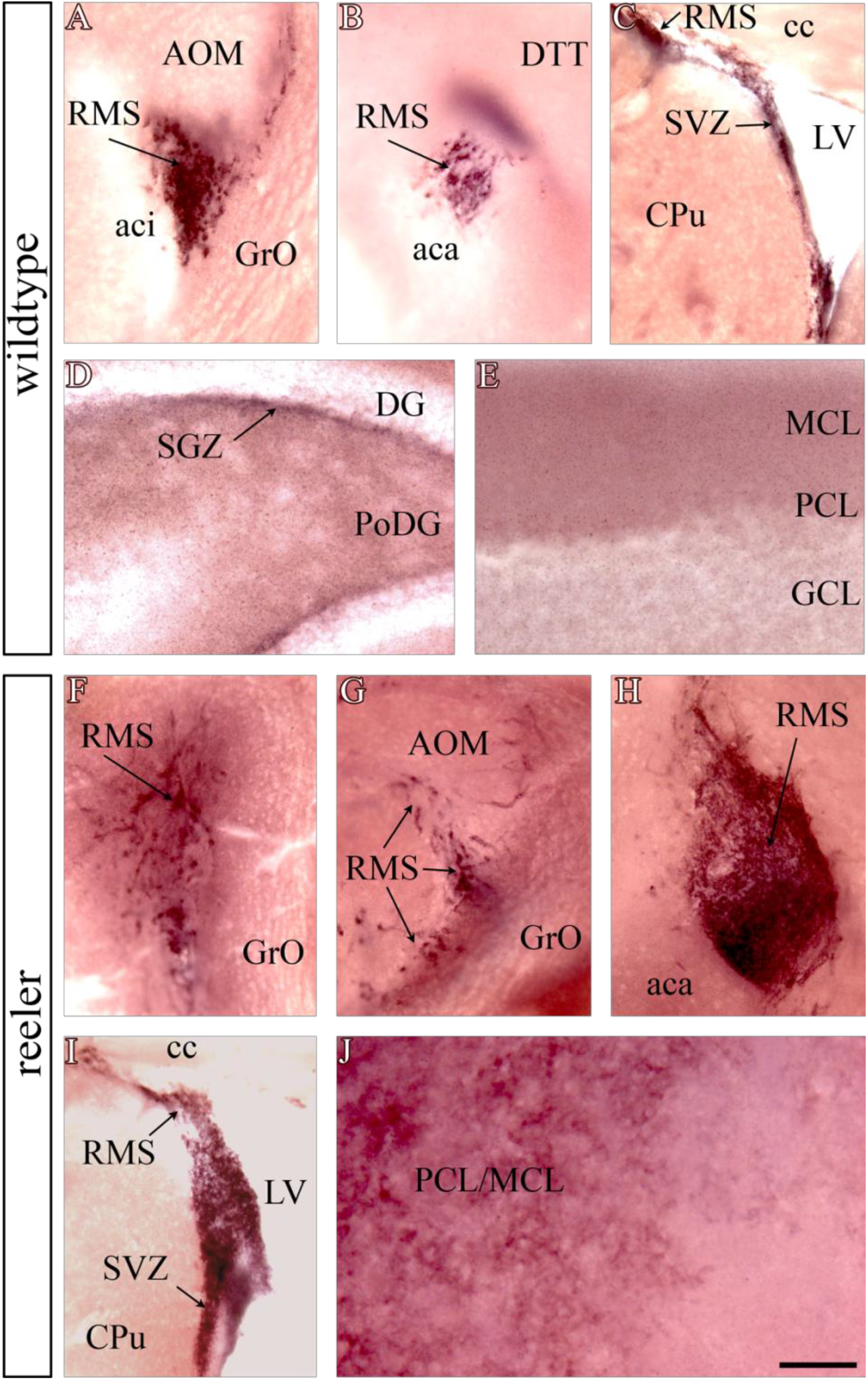
Localization of expression sites of functional reclin receptors in the adult brain. AP-reelin staining in wild-type (A-E) and *reeler* (F-J) mice reveals the presence of functional reelin receptors along the SEZ/RMS in both wild-type (A-C) and *reeler* (F-I). The RMS of *reeler* mutants appears very thin (hypocellular) in its rostral portion (F-G), with no signs of neuroblast accumulation in the nucleus of the olfactory bulb, but considerably thicker (extremely hypercellular) in its caudal part (H-I). FRRs are also found in the subgranular area of the hippocampus (D). Note the weak staining of the molecular layer of the cerebellum in the wild type (E), which is markedly increased in *reeler* (J). aca: anterior part of the anterior commissure; aci: intrabulbar part of the anterior commissure; AOM: medial part of the anterior olfactory nucleus; cc: corpus callosum; CPu: Caudate-Putamen nucleus; DTT: dorsal tenia tecta; DG: dentate gyrus; GCL: cerebellar granular layer; GrO: granular layer of the olfactory bulb; LV: lateral ventricle; MCL: cerebellar molecular layer; PCL: Purkinje cell layer; PoDG: polymorphic layer of the dentate gyrus; RMS: rostral migratory pathway; SVZ: subventricular zone. Scale bar: 200 μm.

### Functional ApoER2 is present in the main cellular compartments of the EZ/SEZ/RMS neurogenic axis

By looking at the specific regions in which ApoER2 is expressed in the brain, the next objective was to determine which cell types of the neurogenic niche of the lateral ventricles would be competent to respond to a signal triggered by reelin, as well as to examine the localization of ApoER2 in the cellular components building up the RMS. The combination of the in situ binding test of the AP-RR36 probe with the detection by immunofluorescence, shows that, to a greater or lesser extent, all the cellular subpopulations of the EZ / SEZ are again competent to receive reelin as a ligand, as demonstrated by the colocalization of functional ApoER2 with specific markers for ependymocytes (E-type cells) [S100β], subventricular astrocytes (including B-type cells) [GFAP], transient amplifying cells (type C cells) [Mash1], neuroblasts (cells type A) [DCX], as well as cells in full S phase of the cell cycle [pulse and hunting after 1 hour of injection with BrdU] (**Figure 2**). It is noteworthy that the highest colocalization in the expression of ApoER2 occurs in the neuroblasts. Therefore, an embryo-adult continuity is expressed in the expression of ApoER2 in the different cell lineages of the neurogenic niche.

**Fig. 2:**
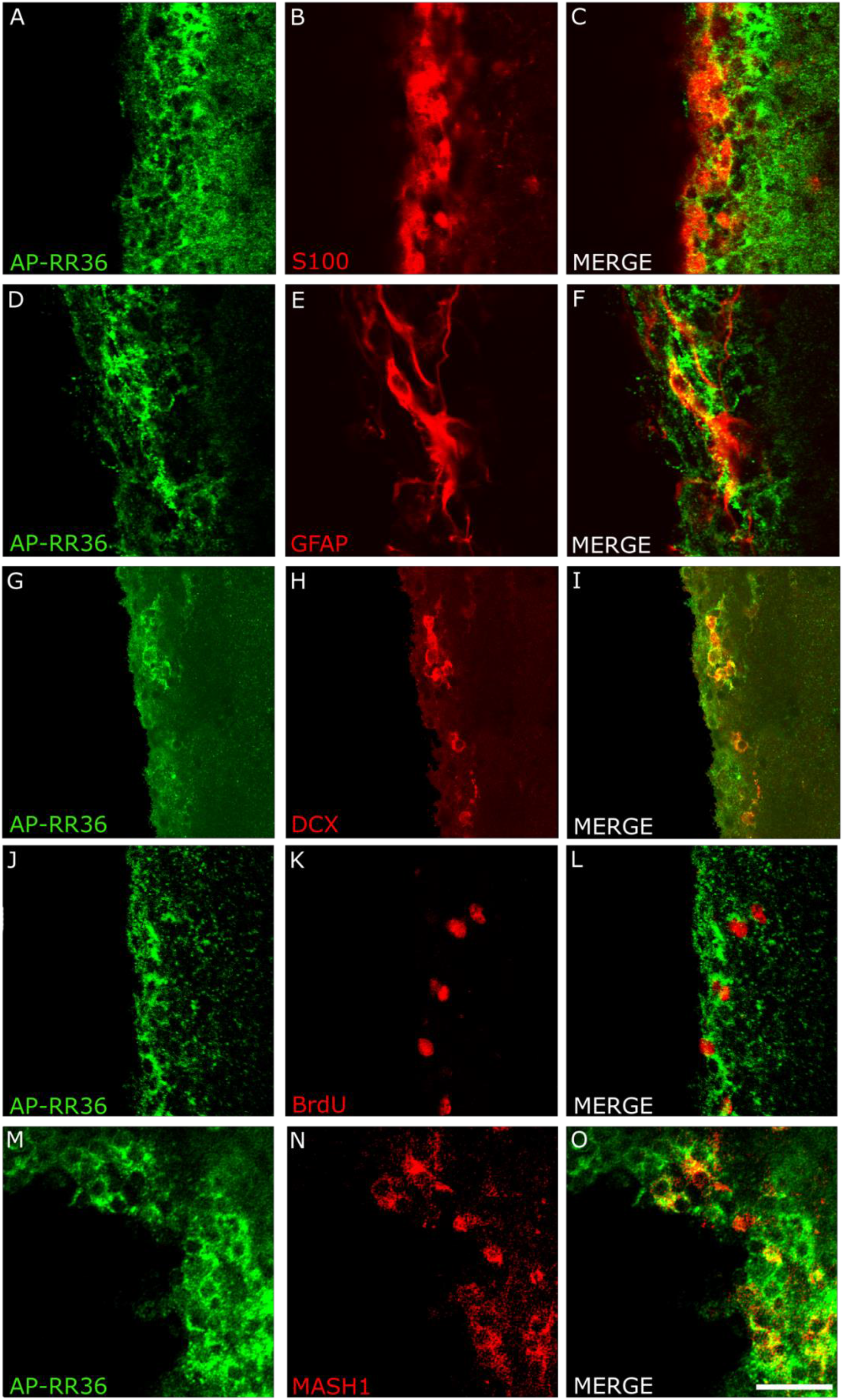
Reelin binds to the main cell types of the major adult neurogenic niche. All EZ/SEZ cell subpopulations are competent to receive reelin as ligand. Colocalization of functional ApoER2 with specific markers for S100b+ cells, corresponding to ependymal cells, type E cells (A-C). Subventricular astrocytes, known as type B cells, which express GFAP, also possess some degree of reelin receptor expression (D-F). The highest degree of identity in expression is observed between ApoER2 and doublecortin, a marker of neuroblasts, or type A cells (G-I). Cells in S phase of the cell cycle (1 hour after BrdU injection, J-L) and so-called transient amplifying cells or type C cells, which express Mash1, are also competent to reelin binding. Scale bar: A-C; G-I: 50 μm. D-E: 30 μm; M-O: 40 μm.

As regards the components of the rostral migratory route, the neuroblasts (positive for the doublecortin marker, DCX) consistently continue to express ApoER2, together with the astrocytes (GFAP +) that make up the “tubes” through which they run and which facilitate its migration through an adult environment. As previously described, neuroblasts also carry out divisions along the RMS, from their birth in the EZ / SEZ until they reach their destination in the different layers of the olfactory bulb (Menezes et al., 1995). It is no less significant that these elements in division would also be competent to respond to reelin (**Supplementary Fig. 3**).

### Reelin signaling remains active in the EZ/SEZ

Although ApoER2 is considered a promiscuous receptor, the indications that until now we had to think that the signaling triggered by reelin was still active in the adult, were more than enough to justify the following experiment.

Our group had observed that the adult reeler mutant showed an exaggerated ventricular enlargement and a hyperplastic RMS. The cells of the EZ / SEZ / RMS are competent to receive the reelin signal, that is, they express one of the canonical elements of the signaling pathway, the ApoER2 receptor. In addition, the presence of the intracellular adapter Dab1 has been described in the RMS, another of the traditional components of the pathway (Hack et al., 2002).

As is known, a condition that must be met for the reelin signal through its receptor to be transduced is the phosphorylation of Dab1 (Howell et al., 1999). During embryonic development, we have shown that reelin, through a mechanism of low regulation of ApoER2, promotes the new neurons dissociate from their parents and is able to normalize the levels of Dab1 on a reeler genetic background (Uchida et al., 2009). With the aim of checking if the reelin signaling pathway is still active in the adult brain, we performed an *ex vivo* Dab1 phosphorylation assay under reelin stimulation, using a homogenate of cells extracted from the EZ as substrate. / SEZ of adult animals reeler.

The reelin fragment comprising the repeats 3 to 6 (RR36) recapitulates the vast majority of the functions of the protein and is capable of triggering a signal (Jossin et al., 2004). In order to continue with the line of experiments carried out so far, for carrying out this phosphorylation assay of Dab1, the AP-RR36 fusion protein was used (although previously and, with the same result, we had already done it with recombinant reelin, generated in our laboratory by transfection of HEK293T cells). Once the incubation time had elapsed, an immunoprecipitation of Dab1 was necessary to enrich the sample with this protein, since an antibody that generically detects the phosphorylated tyrosine residues would be used.

The results of this test show that, in effect, Dab1 phosphorylation occurs after incubation of EZ / SEZ cells with AP-RR36. The increase of the signal of the phosphotyrosine band, together with the decrease of the Dab1 signal, consistent with the beginning of its degradation, allows to affirm that the reelin signaling persists in the adult EZ / SEZ (**Figure 3**).

**Fig. 3:**
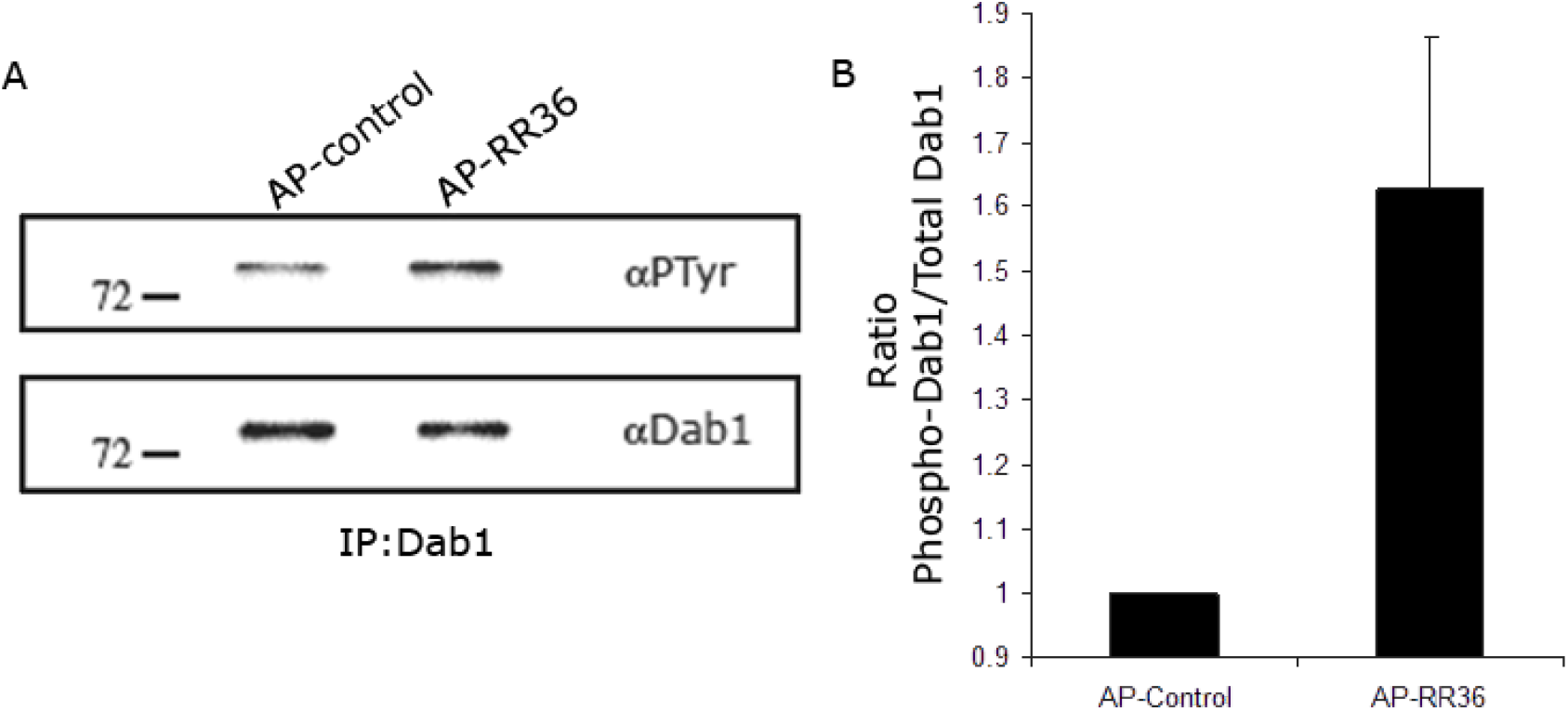
Reelin signaling remains active in the adult SVZ. Dab1 phosphorylation ex vivo assay after stimulation of a dissected tissue cell extract from the EZ/SEZ of reeler brains with the AP-RR36 fusion protein. Due to the lack of specific antibodies to phosphorylated Dab1 tyrosine residues, immunoprecipitation of this protein is necessary to enrich the sample for detection of phosphorylated tyrosine residues by Western blot (A). The addition of the central fragment of reelin causes a 60% increase in the ratio between phosphorylated Dab1 and total Dab1, in parallel with a 20% degradation of the total Dab1 present in the sample (B).

### Lack of function of reelin, ApoER2 or Dab1, but not of VLDLR, causes analogous phenotypes in the lateral ventricles and RMS

The lack of function of reelin, ApoER2 or Dab1 in mutants lacking these proteins, causes hyperplasia of the RMS (Andrade et al., 2007). Closely related to this structure are the lateral ventricles of the adult brain. When analyzing the neuroanatomy of these mutants, we observed that they precisely show an evident ventriculomegaly (increase in ventricular size, Fig. 4B, C, E), but not the VLDLR mutant (Fig. 4D), which does not exhibit phenotypic alteration of any of these structures (which supports that the main receptor of reelin in the neurogenic niches of the adult brain, if not the only one, is ApoER2).

**Fig. 4:**
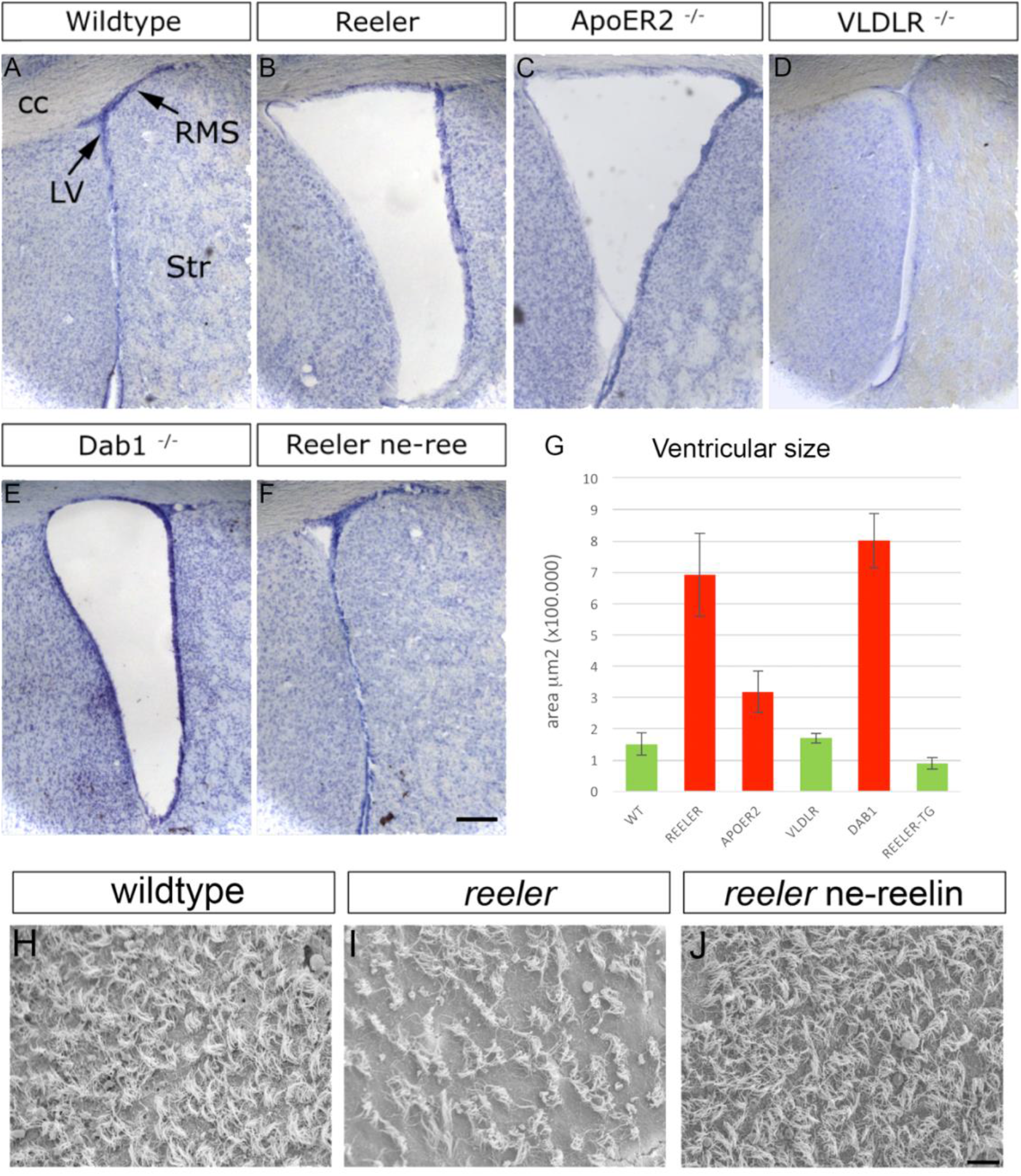
Increased ventricular volume in mutants related to reelin and its signaling pathway and recovery of ependymal cilia density due to ectopic expression of reelin. Nissl staining showing the size of the lateral ventricles of different mutants compared with the wild-type genotype (A). Null mutants for reelin, ApoER2 and Dab1 (B, C, E) and conditional ablation of Dab1 in progenitors (G, H), show an increase in this structure size, whereas gain of reelin function normalizes this defect (F). The VLDLR mutant exhibits a normal ventricular phenotype (D). Photomicrograph obtained by scanning electron microscopy of the ependymal cilia present in the wall of the lateral ventricle (H). Cilia density and morphology is severely affected in reeler (I). Expression of reelin by neural progenitors in the adult subependymal zone results in normalization of this phenotypic defect (J). Scale bar: (A-F: 100 μm; H-J: 10 μm).

### Partial rescue of lateral ventricle integrity and neuroblast migration by ectopic expression of reelin in NPCs of *reeler* mutants

To define the possible reelin function and its signaling pathway in the adult EZ / SEZ system, we carried out a series of studies with genetically manipulated animals. In the same way as we did before (Pérez-Martínez et al., 2012), we used the ne-reelin transgenic mouse to evaluate the effect of the reelin gain function on a wild background and in a rescue paradigm on a reeler (reeler ne-reelin) adult background. The work in which this transgenic mouse was used for the first time (Magdaleno et al., 2002) highlights the rescue of foliation of the cerebellum in reeler ne-reelin animals, with the consequent disappearance of the ataxic phenotype, although they indicate that the cortical lamination apparently not recovered. As a matter of fact, we demonstrated that it does partially occur above an ectopic subplate (Pérez-Martínez et al., 2012). The analysis of the expression of ectopic reelin in the adult, now provides a key information to deepen the understanding of the reelin function.

The protein expressed ectopically by the neural progenitors of the EZ / SEZ (nestin promoter) on a reeler background causes a normalization of the ventricular size (**Fig. 4B, F**) and, in fact, a recovery of the density of ependymal cilia, analyzed by scanning electron microscopy (**Fig. 4, H-J**). The graphic representation of the values obtained by means of software shows the statistically significant differences between the ventricular size of the reeler mutants, Dab1 and ApoER2 (**Fig, 4G**, columns in red).

### Conditional ablation Dab1 in NPCs impairs neuroblast clearance from the SVZ

To address this issue from the opposite side, we decided to carry out a conditional ablation of Dab1, since this protein is expressed in both the embryonic radial glia (Luque et al., 2003) and the adult EZ / SEZ (Hack et al., 2002), which would provide data that would complement perfectly with those we had obtained with gain-of-function experiments. With this purpose, we made use of the CRE-lox system induced by Tamoxifen and thus cause the interruption of reelin signaling in progenitors, since it provides the possibility of selecting the place and the moment in which we wish to eliminate the gene of interest, Dab1 in this case. Through crosses between the line of transgenic mice nestin-CRE^ERT2^ (expression of the enzyme CRE recombinase under the nestin promoter, Balordi and Fishell, 2007) and the floxed-Dab1 line (the loxP targets of the enzyme flank the Dab1 gene; Pramatarova et al., 2008; Teixeira et al., 2012), we provoke the conditional ablation of reelin signaling in the progenitors of adult mice. The effective ablation of Dab1 after the application of Tamoxifen, is evident by immunodetection of the protein in the RMS at the level of the olfactory bulb, as well as in more caudal regions such as in the area of the anterior commissure, the lateral ventricles or the zone. subgranular of the hippocampus, these last places being neurogenic niches where nestin is expressed with more intensity (**Supplementary Fig. 4**). We perform a 3 days BrdU pulse experiment to demonstrate that the conditional ablation of Dab1 in progenitors (**Fig. 5A, B**), produces an increase in the size of the lateral ventricles (**Fig. 5C, D**) and impairs the abandonment of the neuroblasts of the SEZ towards the RMS, accumulating (**Fig. 5E-F**), so it can be affirmed that the phenotype observed in the mutants of the elements of the reelin signaling pathway is not an event that depends on development. Thus, we observed how the interruption of the reelin pathway by the selective elimination of Dab1, results in an accumulation of cells in the more caudal regions of the RMS. Under normal conditions, a neuroblast that is born in the EZ / SEZ takes about 3 days to migrate to the olfactory bulb through the RMS. In this case, in the Dab1flx / flx; CRE + mice, a dramatic increase in BrdU + profiles was observed in the EZ / SEZ, which contrasts with those found in the Dab1 + / flx; CRE + mice, where the great majority of cells that incorporated BrdU have abandoned their neurogenic niche to begin their long journey towards the olfactory bulb. The genetic ablation of Dab1 causes, therefore, a defect in the migration of neuroblasts.

**Fig. 5:**
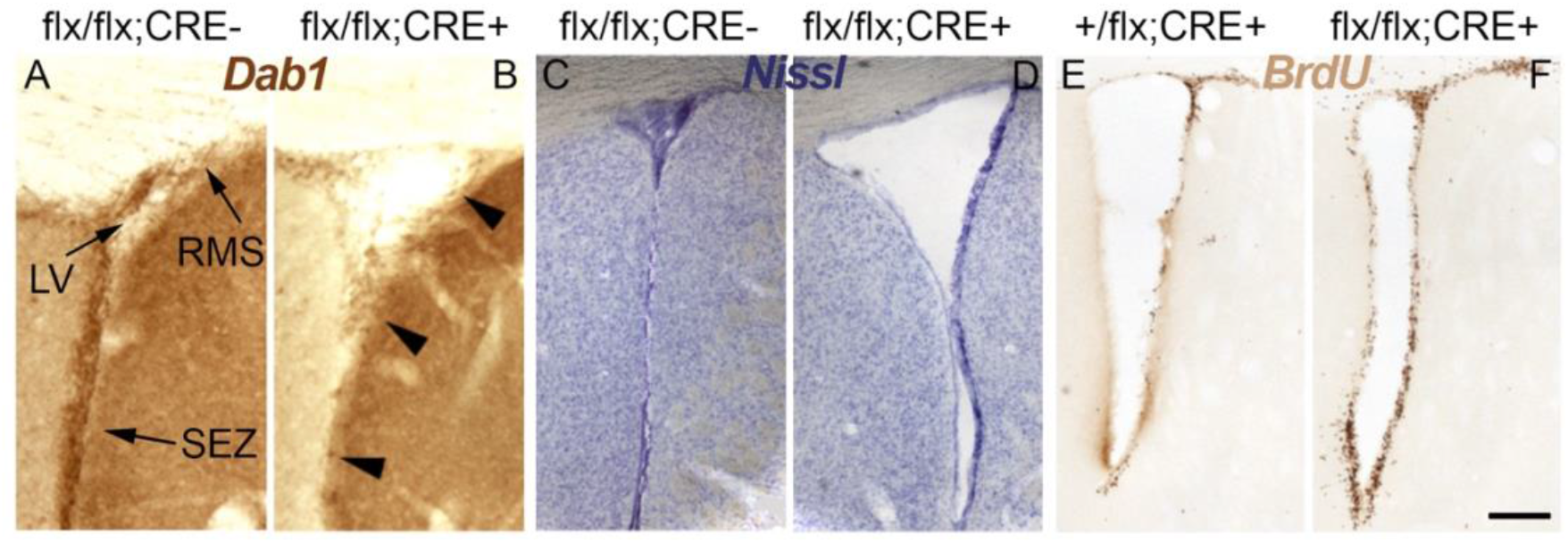
Effects of Dab1 ablation after tamoxifen treatment in adult mice. Immunohistochemical detection of Dab1. The signal is present in neuroblasts of the subependymal zone and SMR (A) and disappears in those cells expressing Cre recombinase enzyme (B). Nissl staining showing enlargement of the lateral ventricle of the Dab1 flx/ flx; CRE+ mouse after exposure to tamoxifen (D), compared to control (C). 3-day BrdU pulse and chase (E, F). In mice with both Dab1 floxed alleles (F), an accumulation of BrdU+ cells (neuroblasts) is observed along the subventricular zone and in the corticostriatal sulcus, onset of RMS, compared to control (E), where neuroblasts have already left the zone to migrate into the olfactory bulb. Scale bar: 100 μm. [CPu: caudate-putamen nucleus; LV: lateral ventricle; SEZ: subependymal zone; RMS: rostral migratory stream].

In a complementary approach, we found the opposite effect in the transgenic reeler ne-reelin brains. We first tested the function of the ne-reelin transgene in the adult brain by immunofluroescence to detect the presence of reelin in the EZ/SEZ of the lateral ventricle, confirming its expression in this area ectopically (**Fig. 6A-I**). Then, we demonstrated that the accumulation of neuroblasts in the RMS observed in a reeler, with the consequent migratory difficulty to reach the olfactory bulb, tends to normalize in the presence of ectopic reelin in the system, recovering also the morphology of the RMS at the level of the olfactory bulb (**Fig. 6J-L**).

**Fig. 6:**
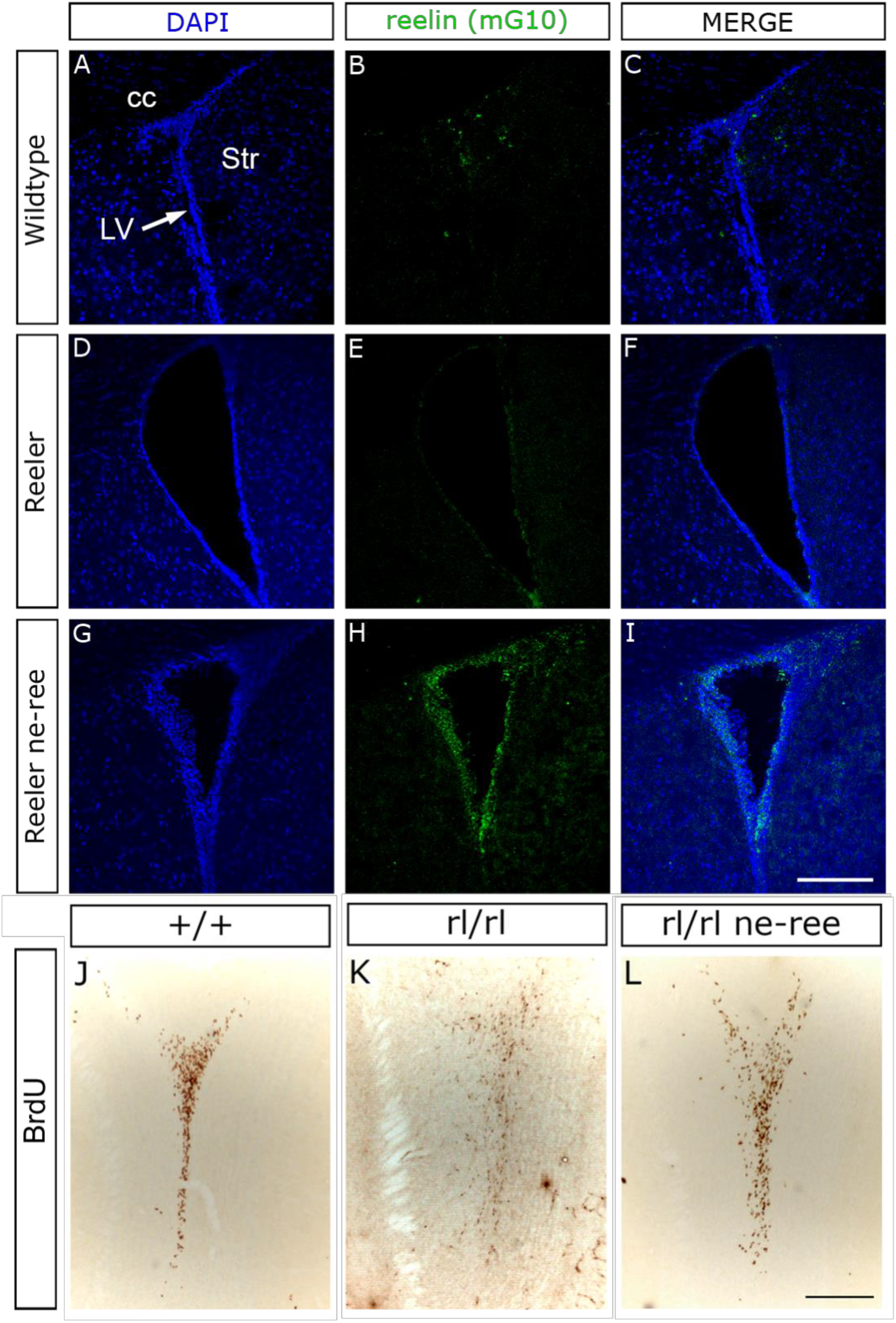
Ectopic reelin expression in the adult nestin-reelin transgenic mouse and rescue of RMS morphology in the nucleus of the olfactory bulb. Immunofluorescence for the detection of reelin using the monoclonal antibody G10. No immunoreactivity to reelin is observed in the wild-type animal (A-C) or in the reeler mutant (D-F). However, the nestin promoter directs reelin expression in the subependymal zone, ectopically, in the reeler ne-reelin transgenic mouse (G-I). Immunohistochemical detection of BrdU after a 3-day pulse and chase experiment in the Wild-type phenotype (J). In reeler, cells appear scattered and in less quantity compared to control (K). The presence of ectopic reelin on reeler background rescues this phenotype, both in morphology and in the number of cells reaching the center of the olfactory bulb (L). [cc: corpus callosum; LV: lateral ventricle; Str: striate nucleus]. Scale bar: A-I: 150 μm; J-L: 200 μm.

### Intraventricular infusion of reelin alters the migration of neuroblasts

Taken together, the previous results point to a reelin action in the maintenance of adult neurogenic niches. The interruption of the reelin signaling pathway causes defects in the migration of the neuroblasts, together with an alteration in the anatomy of the lateral ventricles. In a further complementary approach, in an environment with excess reelin, we used osmotic mini-pumps to administer recombinant reelin directly into the lateral ventricle of adult reeler mice (**Fig. 7A**). The pumps were carefully implanted using stereotaxic surgery and left to work for 5 days, releasing recombinant reelin at a rate of 1μl / hour. Subsequently, immunohistochemical techniques were performed to detect Doublecortin + cells (neuroblasts). The infusion of control culture medium does not alter the distribution of the neuroblasts (**Fig. 7B**), whereas the infusion of reelin affects the migratory behavior of the neuroblasts on a reeler genetic background, producing a remarkable alteration in the usual location of the neuroblasts on the side ipsilateral to that of the cannula insertion of the reelin pump (**Fig. 7C**, arrowheads in red). Therefore, we can conclude that reelin influences the migration of neuroblasts in the adult brain, and probably does it from the CSF, which in fact, as our group showed before, naturally contains it (Sáez-Valero et al., 2004).

**Fig. 7:**
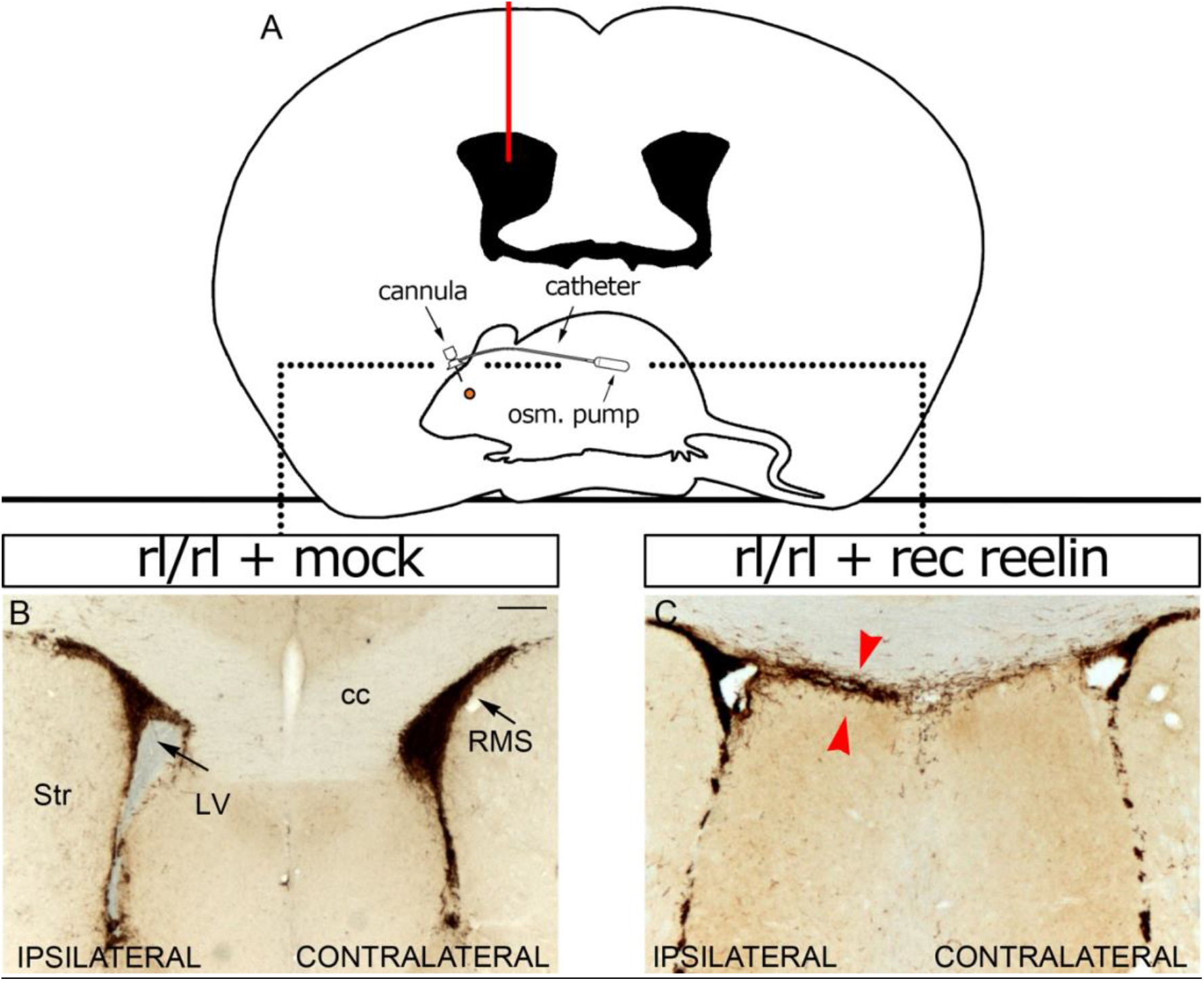
Reelin modifies neuroblast migration in vivo. Infusion of recombinant reelin or control medium for 5 days into the right ventricle of adult reeler mice by implantation of osmotic mini-pumps (A). Infusion of control medium produces no change in neuroblast distribution, as evidenced by this immunohistochemistry for detection of doublecortin (B). In contrast, the presence of recombinant reelin dramatically alters the normal localization of neuroblasts around the ventricle ipsilateral to the infusion (C, arrowheads). [cc: corpus callosum; LV: lateral ventricle; RMS: rostral migratory pathway; Str: striatal nucleus]. Scale bar: 200 μm.

## Discussion

Here we show that in the adult brain the reelin-activated biochemical-signaling cascade, resulting in phosphorylation of the intracellular adapter Dab1, occurs in the ependymal-subependymal zone (EZ/SVZ) of the lateral ventricles. We also show that the EZ/SEZ, localizing stem cells and other neural progenitors, and its associated and extensive rostral migratory stream (RMS) of neuroblasts, concentrate the highest density of functional ApoER2 in the adult brain. Mutant mice lacking reelin (*reeler*), ApoER2 or Dab1exhibit a prominent widening of ventricular volume and a marked hypercellularity of the RMS, particularly in its proximal aspect. The conditional ablation of Dab1 in neural progenitor cells of wild types increases the ventricular volume and inhibits cell migration from the SVZ to the RMS. In contrast, the ectopic transgene expression of reelin in the neural progenitors of *reeler* mutants normalizes the ventricular lumen and the density of ependymal cilia, improving in turn the migration of neuroblasts; consistently, the intraventricular infusion of recombinant reelin in *reeler* mutants also affects the migration of neuroblasts, indicating that reelin acts from the cerebrospinal fluid. These results show that reelin signaling not only remains active, but also is necessary and sufficient to modulate the adult EZ/SEZ/RMS neurogenic axis.

The dynamics and molecular composition of cerebrospinal fluid (CSF) are the object of renewed interest, particularly insofar as it also emerges as a key component in the regulation of the neurogenic niche EZ/SVZ (reviewed in Lun et al., 2015). The presence of reelin in the CSF of the adult brain has been documented by our group (Sáez-Valero et al., 2003) and later confirmed by others (eg, Ignatova et al., 2004, Guldbrandsen et al., 2014; Macron et al. al., 2018). Various mechanisms are involved in the complex dynamics of the CSF (reviewed in Iliff et al., 2013; Hladky and Barrand, 2014). One of them, the orchestrated movement of the ependymal cilia of the ventricular wall causes a laminar supraependimal flow of approximately 200 μm in thickness (Worthington and Cathcart, 1966, Cifuentes et al., 1994, Siyahhan et al., 2014). It has been observed that this type of movement is not only necessary for the normal flow of CSF in the lateral ventricle, but also for the formation of gradients of concentration of molecules contained in the fluid and for the directional migration of the neuroblasts that runs parallel to said flow (Sawamoto et al., 2006). In addition, multiple evidences relate the ciliary alterations and the planar polarity of the ependymocytes with the ventricular distension and the hydrocephalus (reviewed in Jiménez et al., 2014; Ohata and Alvarez-Buylla, 2016). Our results involving the activity of reelin in maintaining the integrity of the ciliated mantle ependymal / ventricular volume and the regulation of neurogenesis/migration of neuroblasts are consistent with these observations. On the other hand, the small apical surface of the B cells is in direct contact with the CSF whose factors in solution can modulate the behavior of these stem parents (Lehtinen et al., 2011; Zappaterra et al., 2007). Moreover, as radial glia and neuroepithelial cells do during embryonic development, the vast majority of B cells contact the ventricle through small apical processes containing a single primary cilium (Mirzadeh et al., 2008; Shen et al., 2008). This primary cilium could well integrate signals directly from molecules present in the CSF (reviewed in Fuentealba et al., 2012). In another vein, the ependymal cells themselves (which surround the apical processes of the B cells), the main interface between the CSF and the brain parenchyma, are linked together by “gap junctions”, a type of intercellular connection that does not it provides a severe restriction to the diffusion of molecules between both compartments (Whish et al., 2015). Undoubtedly intriguing is the detection of ApoER2 and VLDLR (as well as that of Notch and some of its ligands) in the cerebrospinal fluid itself (Guldbrandsen et al., 2014; Macron et al., 2018). Perhaps its presence may be related to the release of extracellular membrane particles (prominin/CD133+) from neural progenitor cells and other epithelial cells to CSF (Marzesco et al., 2005; Huttner et al., 2008). Where does the CSF reelin come from then? Most likely, CSF reelin is not derived from blood but from the CNS (Ignatova et al., 2004, Aasebø et al., 2014). On the other hand, although its expression is detectable during embryonic development (Lein et al., 2007; Johansson et al., 2013, 2014), reelin was not found in the secretome of the choroidal plexus of the lateral ventricle, classically considered a primary producer of CSF, and recently revealed as a key component of the adult EZ/SVZ niche (Silva-Vargas et al., 2016). However, it is possible that reelin reaches the neurogenic niche EZ/SVZ after being secreted into the CSF from some other circumventricular organ (CVO). For example, the subfornical organ, located on the ventral surface of the fornix near the foramen of Monro that interconnects the lateral ventricles with the third ventricle, strongly expresses reelin (Lein et al., 2007). Traditionally considered a sensory CVO, its secretory capacity has recently been revealed (Agassandian et al., 2017). It also expresses reelina the subcomisural organ (SCO), located in the dorsocaudal region of the third ventricle, at the entrance of the cerebral aqueduct (Lein et al., 2007). It is known that the SCO secretes SCO-spondine, transtirretin and basic fibroblast growth factor (BFGF), proteins that participate in various aspects of neurogenesis such as the proliferation of stem progenitors and neuronal differentiation, including axonal guidance. Some unidentified soluble compounds secreted by the SCO have been detected in the lateral ventricle (Guerra et al., 2015). In structural terms, the main SCO secretion, SCO-spondine, belongs to the TSR superfamily (thrombospondin type 1 repeat), which includes F-spondine and thrombospondin-1 among other proteins (Adams and Tucker, 2000). Reelin possesses a domain homologous to the N-terminus of F-spondine (Ranaivoson et al., 2015) and in turn the F-spondine protein is able to interact, through its TSR domains, with ApoER2 (Hoe et al., 2005). Thrombospondin-1 (TBHS-1) can interact with ApoER2 and VLDLR and it has been proposed that TBHS-1 stabilizes the chains of neural precursors derived from explants of the SVZ (Blake et al., 2008), however, its function *in vivo* remains to be clarified. Interestingly, the lack of secretory function of the SCO is a common feature in various animal models of hydrocephalus (Meiniel, 2007), classically defined as a distension of the ventricular system resulting from an active accumulation of CFS. Multiple evidences associate the hydrocephalus with abnormal neurogenesis; the most recent results involve neurogenic alterations rather than the active accumulation of CSF in its pathogenesis (Furey et al., 2018). In any case, and independently of where the CSF reelin of the adult brain comes from, there is a suggestive parallelism between this causal association and our own results indicating that the inhibition of reelin activity on the EZ / SVZ niche causes alterations that compromise both the neurogenesis as the ventricular volume.

It is well known that the activation of Notch signaling promotes and maintains the character of neural progenitor, inhibiting neuronal differentiation. The Numb protein, on the other hand, antagonizes the function of Notch during the division of neural precursors (reviewed by Roegiers and Jan, 2004, Engler et al., 2018). The canonical signaling of Notch is in fact very active in the neurogenic niche EZ / SVZ regulating the maintenance of the stem neural progenitors (Imayoshi et al., 2010, Aguirre et al., 2010). Although the precise mechanisms of regulation of the activity of Notch signaling in the mouse EZ-SVZ are not known, it is known that some Notch ligands are expressed along the same and both Delta1 and Jagged1 have been detected in IPCs and also in neuroblasts (Aguirre et al., 2010, Irvin et al., 2004, reviewed in Fuentealba et al., 2012). Interestingly, the postnatal ablation of Numb / Numblike proteins in the EZ-SVZ causes widening of the lateral ventricles as well as a marked hypercellularity of MSY (Kuo et al., 2006), phenotypes coinciding with those described here for the deficiencies of reelina, ApoER2 or Dab1. Consistently, overexpression of the activated intracellular domain of Notch in the EZ-SVZ phenocopy the hypercellularity of the RMS; however, the integrity and size of the lateral ventricles are not compromised, suggesting the possibility that Numb regulation of ependymal integrity occurs through a mechanism independent of Notch (Kuo et al., 2006). In contrast, a more recent study shows how the ependymal cells are dependent on the canonical signaling of Notch, which actively maintains the quiescence, phenotype and position of the same. The inhibition of Notch signaling exclusively in the ventricular ependyma allows the reentry of ependymocytes in the cell cycle and the production of neurons of the olfactory bulb, while the forced signaling of Notch is sufficient to block the plastic response of the ependymal cells to cerebral infarction (Carlén et al., 2009). Undoubtedly, Notch signaling has different functions in the different cellular populations of the neurogenic niche of the lateral ventricle. On the other hand, although the interaction between reelin and Notch signaling is important for the modulation of neurogenesis and cell migration during cortical development (Hashimoto-Torii et al., 2008; Lakoma et al., 2011), molecular mechanisms precise have not yet been clarified. In fact, a hundred of genes are directly regulated *in vivo* in cortical NSCs by the intracellular domain of Notch (NICD) and its co-factor RPBJ. One of them is Dab1, whose transcription after the activation of Notch, is not activated, but repressed (Li et al., 2012). Moreover, Numb and Dab1 are endocytic accessory proteins, capable of regulating the endocytosis and the subsequent post-endocytic trafficking of their related receptors [such as Notch, TrkB, b-APP, VLDLR and ApoER2] (Yap and Winckler, 2015). Since both proteins share the PTB domain (Phosphotyrosine binding/NPXY binding region), the possibility of a cross interaction and depending on the cellular context is not ruled out. A recent study demonstrates how the inhibition of Notch signaling in the adult neurogenic niche of the lateral ventricle results in the cells leaving the SVZ migrating towards the RMS more rapidly than that observed in the control brains. In contrast, the overexpression of Notch results in a greater proportion of cells being retained in the SVZ for longer than that observed in the control brains (Piccin et al., 2013), a phenotype clearly similar to that described for conditional ablation of Dab1 in this work. Also, the abundance of Notch in the neuroblasts (PSA-NCAM+) located in the RMS, much lower in those that have reached the OB, suggests that the activity of Notch can inhibit its premature differentiation (Givogri et al., 2006). Again, a pattern of expression strongly reminiscent of the here shown for the localization of ApoER2. Doubtless, it would be interesting to explore if the interaction between reelin and Notch signaling persists in the EZ/SVZ/RMS neurogenic axis of the adult brain.

The literature results of the manipulation of the PSA-NCAM glycoconjugate are also of potential interest in our context. The poly-*α*2,8-sialic acid (PSA), a unique post-translational modification of the neural cell adhesion molecule (NCAM), is strongly linked to the development and neural plasticity. The elimination of PSA (by means of the genetic ablation of its synthesizing enzymes), independently of NCAM, results in an expansion of the RMS and a ventricular widening (Weinhold et al., 2005), phenotypes comparable to those shown here for the absence of reelina, ApoER2 or Dab1. Also, intrathecal administration of endoneuraminidase (endo N) [an enzyme that selectively eliminates PSA] demonstrates that PSA not only promotes rostral tangential migration of neuroblasts but also suppresses the differentiation thereof induced by NCAM. Said differentiation, dependent on cell-cell contact, involves the activation of the mitogen-activated protein kinase (MAPK) pathway (Petridis et al., 2004). In fact, as in our case ApoER2, in the wild mice the low-regulation of PSA is evident once the neuroblasts reach the OB to begin to migrate radially and differentiate. Interestingly, it has been shown that reelin induces the separation of neuroblasts by migrating in chain from explants of the SVZ (Hack et al., 2002) and that said effect also depends on the activation of the MAPK pathway (Simó et al., 2007). In addition, the intraventricular administration of endo N (and not the elimination of PSA by genetic manipulation) causes a massive dispersion of neuroblasts in surrounding brain areas, including the striatal region (Battista and Rutishauser, 2010); an effect certainly comparable to the redirection of neuroblasts observed here after the intraventricular infusion of recombinant reelin in reeler mutants. There is not known direct interaction between reelin or ApoER2, with PSA or NCAM. However, PSA on NCAM is capable of directly binding to bioactive molecules such as neurotrophins, neurotransmitters and growth factors, involved in a wide variety of neural functions, thus regulating their extracellular concentrations and signaling modes. In this way, PSA binds to the brain-derived neurotrophic factor (BDNF) and BDNF in the PSA chain can migrate to the TrkB and p75NTR receptors (Sato et al., 2016). In turn, neurotrophins, including BDNF, are able to regulate the proteolytic processing of ApoER2 by activating the Trk signaling pathway (Larios et al., 2014). On the other hand, NCAM can interact heterophilically with other cell adhesion molecules such as L1 (Horstkorte et al., 1993). A recent study demonstrates the proteolytic cleavage of L1 by reelin, further indicating its relevance for neuronal migration during cortical development (Lutz et al., 2017). It would be equally interesting to explore if these transitive molecular interactions persist in the EZ/SEZ/RMS neurogenic axis.

Recently it was shown that ependymal and adult neural stem cell share common embryonic progenitors. Number of cells in the lateral ventricle decrease despite ventricle growth and ependymal cells expand apical surface area to accommodate larger ventricle size (Redmond et al., 2019). An impairment of such a mechanism with defective reelin signaling might underlie the ventriculomegaly here observed in reelin, ApoER2 and Dab1 deficient mice. Moreover, reelin signals through mTOR (mammalian target of rapamycin) to regulate dendritic growth (Jossin and Goffinet, 2007). Interestingly, mTORC1 and primary cilia are required for brain ventricle morphogenesis (Foerster et all., 2017), and timing of mTOR activation is also critical for EZ/SEZ/RMS neurogenesis (Magri et al., 2013).

We propose that, acting through ApoER2 and Dab1, CSF reelin controls the integrity and functionality of the ventricular ependymal, and the behavior of the cellular progenitors in the EZ/SEZ; finally regulating the production, differentiation and migration of the RMS neuroblasts at least up to the OB core. While admittedly less obvious, this model is not ad odds with the one we previously proposed for the embryonic development of the cerebral cortex (Pérez-Martínez et al., 2012). In fact, the requirement for a down-regulation of ApoER2 in the neuroblasts along the RMS in the wild mice is clearly deductible from the higher intensity of functional ApoER2 observed in the *reeler* mutant. Taking into account that the neuroblasts while transiting the RMS have not left the cell cycle, that is, they do not constitute an entirely post-mitotic cellular population, in no way ApoER2 down-regulation would reach the minimum observed in the embryonic cortical plate. However, once the neuroblasts reach the core of the OB, a complete down-regulation of ApoER2 could be required to enable (perhaps by signals other than reelin) the radial migration of the olfactory interneurons produced in the adult neurogenic niche. Needless to say, the consistent detection of functional ApoER2 in the neurogenic niche of the subgranular zone of the adult hippocampal dentate gyrus support our notion and deserves further investigation. We believe our results open new avenues of research foreshadowing reelin signaling as an integral part of a master mechanism coupling neurogenesis and neuronal migration for life.

## Acknowledgements

This work was supported by grants from the Spanish Ministry of Science and Innovation SAF2004-07685 and the Fundación Médica Mutua Madrileña [to JML]. We deeply appreciate Dr. J. Nimpf for ApoR2- and Vldlr-null mice; Dr. B. Howell for Dab1-null and conditional (floxed) Dab1 (Dab1c/c) mice, and for Dab1 (B3) antibody; Dr. M. Hattori for the AP-RR36 reelin fusion protein. We also thank Dr. T. Curran for the Nestin-reelin allele and the reelin pCrl plasmid, and Dr. G. Fishell for the Nestin-CreER allele. To our fathers*, in memoriam*.

## SUPPLEMENTARY FIGURES

**Fig. S1.**
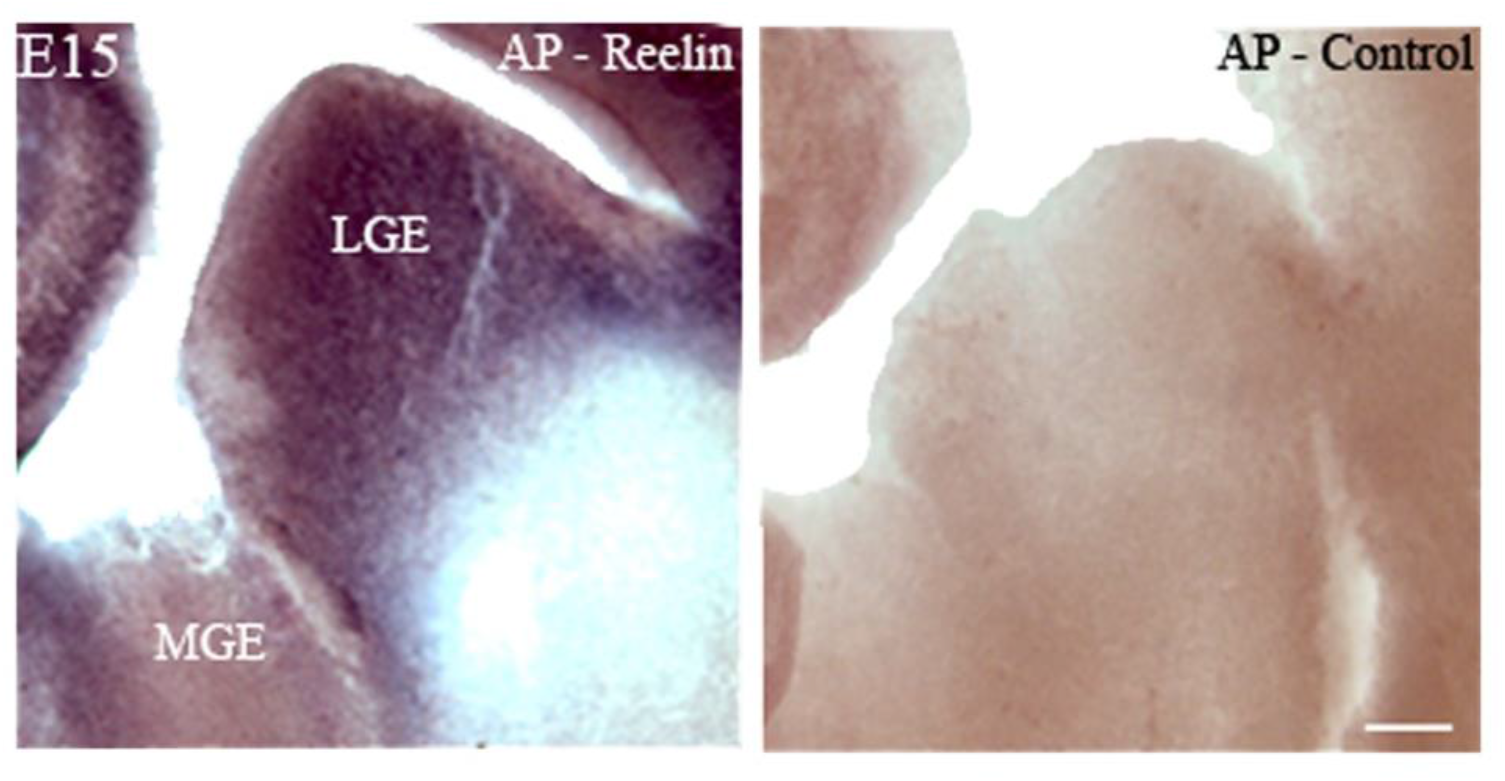
Functional Reelin receptors are expressed in the LGE. In situ binding with the AP-RR36 fusion protein and AP-Control on a mouse brain section at day 15 of embryonic development. The staining, corresponding to functional Reelin receptors, is circumscribed to the lateral ganglionic eminence (LGE), embryonic origin of the adult EZ/SVZ/RMS, leaving the medial ganglionic eminence (MGE) free of labeling. Scale bar: 100 μm.

**Fig. S2:**
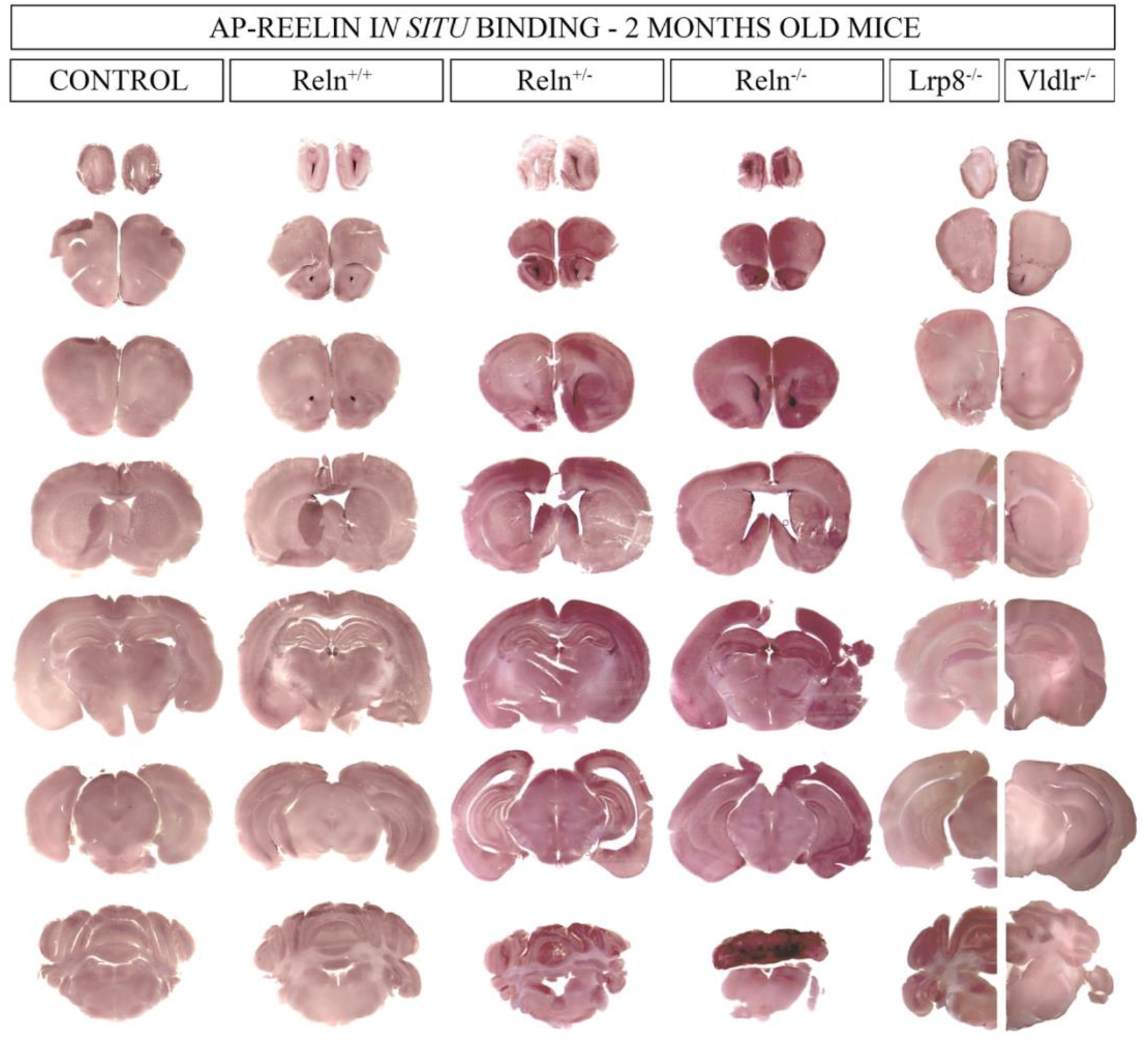
The major reelin receptor in the neurogenic niches of the adult mouse brain is ApoER2. Expression mapping of reelin binding sites shown along the rostro-caudal axis, from the olfactory bulb to the cerebellum, revealed using the AP-RR36 fusion protein. The site of maximal expression is located in the rostral migratory pathway, the subventricular zone of the lateral ventricles, the subgranular layer of the hippocampus, and the molecular layer of the cerebellum. Note an increase in signal as the gene dose of reelin decreases, reaching a maximum level in *reeler* (Reln-/-). The Lrp8 mutant, which lacks the ApoER2 receptor, shows virtually only signal in the cerebellum, indicating that it is the main, if not the only, reelin receptor in the neurogenic niches of the adult brain, in contrast to the VLDLR mutant, which exhibits virtually wild-type staining throughout the brain, except in the cerebellum, where a decrease in signal is seen, indicating that VLDLR is the main receptor in this structure.

**Fig. S3:**
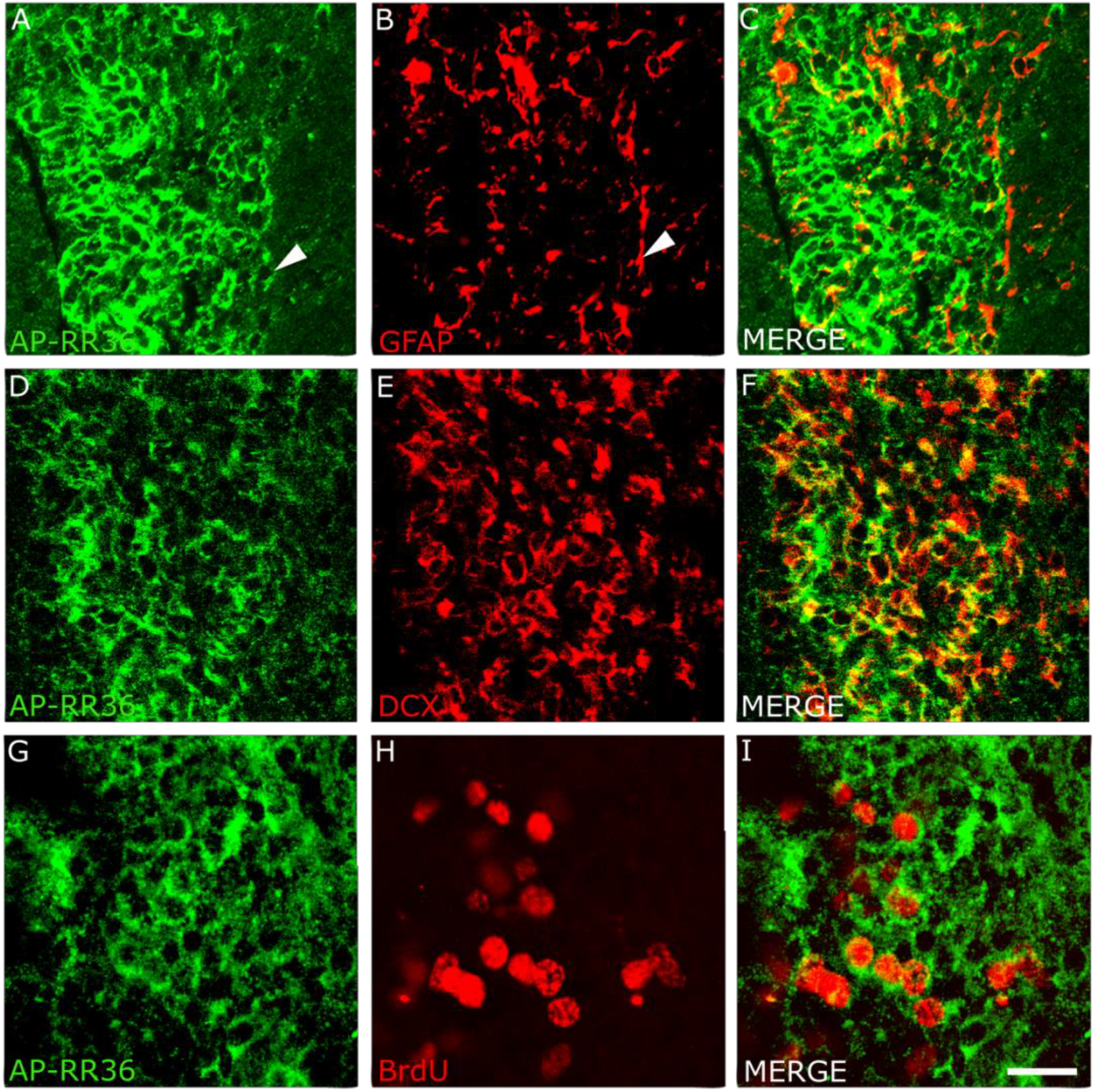
Functional Reelin Receptors are expressed in the RMS. The expression of Functional Reelin Receptors is maintained along the rostral migratory pathway, as shown by this colocalization of the AP-RR36 fusion protein and GFAP+ atrocytes (A-C), which form the tubes through which the neuroblasts, DCX+ (D-F), pass. Cells in S phase of the cell cycle, located along the entire RMS, are also competent to Reelin action (G-H). Scale bar: A-F: 40 μm; G-I. 30 μm.

**Fig. S4:**
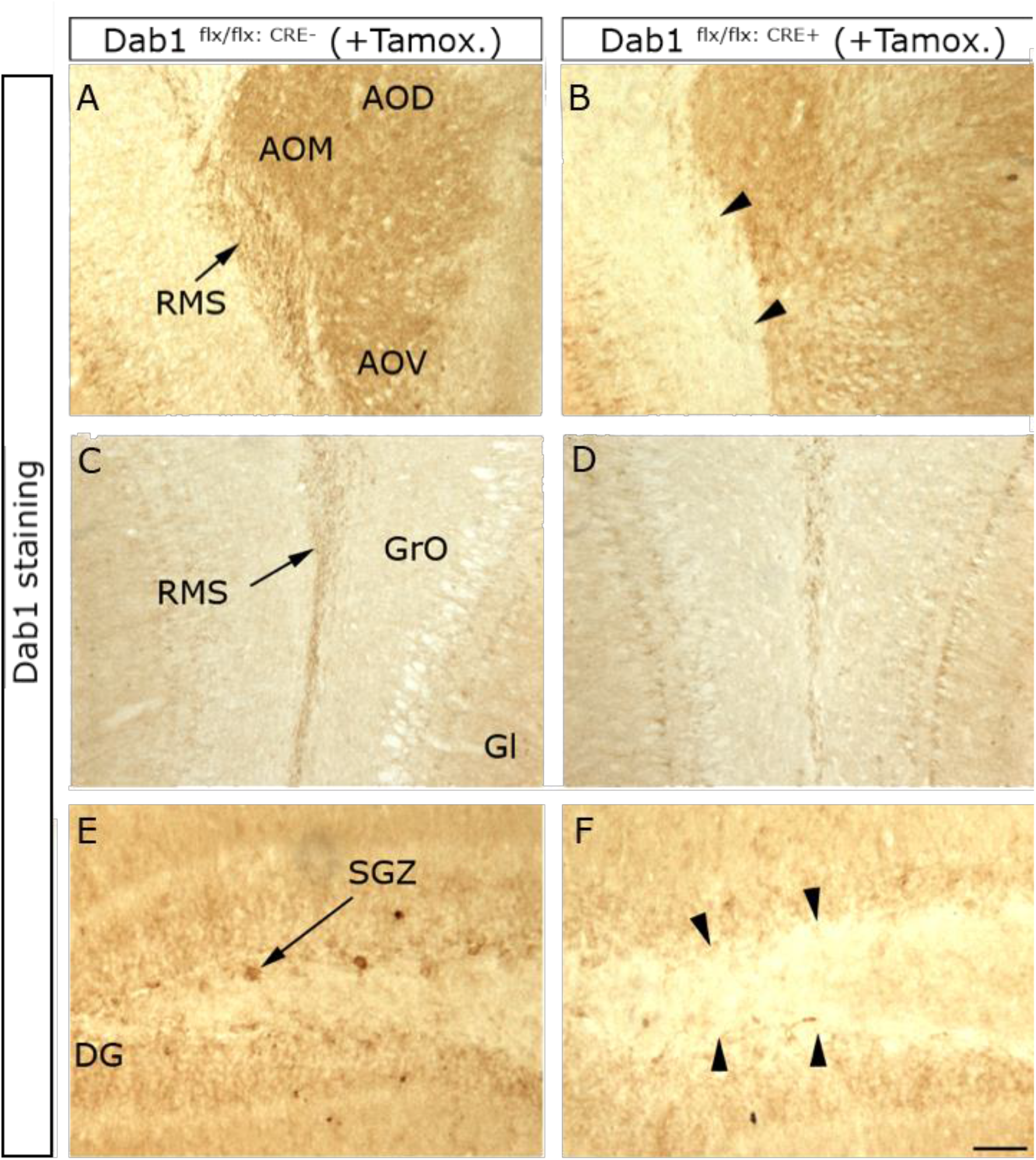
Conditional ablation of Dab1. Immunodetection of Dab1 after 8 days of tamoxifen treatment in the inducible CRE-lox system. CRE recombinase is expressed in this case in neural progenitors under the nestin promoter. The Dab1 signal observed in the RMS at the level of the anterior olfactory nucleus (A, arrow) is absent in the CRE+ individual (B, arrowheads). In the center of the olfactory bulb, Dab1 signal is detected in both control (C) and CRE+ animals (D), consistent with the pattern of nestin expression. In the hippocampal subgranular zone (SGZ), where neurogenic activity persists, Dab1-positive cellular profiles (E) disappear in the presence of recombinase. Scale bar: AB: 100 μm; CD: 200 μm; EF: 50 μm. [DG: dentate gyrus; AOM: medial part of the anterior olfactory nucleus, AOD: dorsal: AOV: ventral; GrO: granular layer of the olfactory bulb; Gl: glomerular layer of the olfactory bulb].

## Notes

### Competing Interest Statement

The authors have declared no competing interest.

